# Calcium signals at cytoneme contacts amplify Wnt/β- catenin signalling

**DOI:** 10.64898/2026.09.15.751720

**Authors:** Emma Cooper, Corin Liddle, Steffen Scholpp

## Abstract

Wnt signalling controls morphogenesis, patterning and homeostasis across tissues. Because Wnt ligands are lipid-modified, their movement between cells cannot rely on simple diffusion and instead requires specialised transport mechanisms. Signalling filopodia, also known as cytonemes, have emerged as a key mediator of contact-dependent morphogen delivery. The mechanisms that stabilise signalling interfaces between Wnt ligand-bearing and receptor-bearing membranes at the cytoneme contact sites, a prerequisite for efficient Wnt ligand-receptor complex assembly and downstream pathway activation, remain poorly understood.

Here, we show that HeLa cell cytonemes carry canonical Wnt components and form contact sites marked by focal Ca^2⁺^ transients. At these contacts, local Wnt and Ca^2⁺^ signals act in a positive feedback loop to promote longer-lasting interactions between neighbouring cells. Mechanistically, we find that E-cadherin/Cadherin 1 (CDH1) co-accumulates at these cytoneme contact sites in a Ca^2⁺^- and Wnt-dependent manner, suggesting that contact stabilisation can be mediated by local adhesion. Consistent with this, presentation of Wnt ligands and CDH1 at cytoneme tips, together with Ca^2⁺^ availability, enhances Wnt signalosome formation at contact sites and, consequently, nuclear β-catenin accumulation in the target cells.

Our data support a model in which local Ca^2⁺^ transients stabilise cytoneme contacts and facilitate efficient Wnt ligand transfer to the receiving membrane, thereby enhancing canonical Wnt/β-catenin signalling. We propose that focal Ca^2⁺^ signalling can serve as a contact-authentication step for morphogen signalling, converting transient cytoneme encounters into effective activators of the signal transduction cascade.

## Introduction

Wnt signalling controls cell fate specification, proliferation, migration, polarity and tissue morphogenesis during embryonic development, and its dysregulation contributes to developmental disorders, fibrosis and cancer (Nusse and Clevers, 2017; Rim et al., 2022). Wnt outputs are commonly grouped into three branches: canonical Wnt/β-catenin signalling and the non-canonical or β-catenin-independent cascades: planar cell polarity (PCP) signalling and Wnt/Ca^2⁺^ signalling (Komiya and Habas, 2008; Niehrs, 2012). However, the classification of the Wnt signalling network does not always predict pathway output in a given tissue. Wnt activity depends on ligand identity, receptor and co-receptor composition, membrane context and cell state, such that the same ligand can elicit distinct downstream responses in different settings (Mikels and Nusse, 2006; van Amerongen and Nusse, 2009). Furthermore, the multiple interconnected branches have traditionally been viewed as functionally antagonistic. For example, the relationship between Wnt/β-catenin and Wnt/Ca^2⁺^ signalling remains unclear, particularly how and when local Ca^2⁺^ signals intersect with canonical Wnt reception at cell–cell contacts (Niehrs, 2012).

Wnt ligands need to move between cells to execute their paracrine functions; however, their mode of transport remains a subject of ongoing debate. Wnts are lipid-modified by mono-palmitoleoylation, and this strongly limits unrestricted diffusion through the aqueous extracellular space (Routledge and Scholpp, 2019; Takada et al., 2006; Willert et al., 2003). Increasing evidence indicates that cytonemes provide a mechanism for spatially controlled morphogen spreading. Cytonemes are thin, actin-rich signalling filopodia that form directed membrane contacts between cells and support contact-dependent transfer of morphogens (Kornberg and Roy, 2014; Ramírez-Weber and Kornberg, 1999; Zhang and Scholpp, 2019). Wnt/β-catenin morphogens can be transported on cytonemes during zebrafish gastrulation to pattern the neural plate (Brunt et al., 2021; Stanganello et al., 2015) and to regulate cancer cell proliferation (Routledge et al., 2022). In addition to ligands, active Wnt ligand-receptor complexes, such as Wnt5b/Ror2, can also move between cells along filopodia in the zebrafish gastrula and in cancer-associated fibroblasts, supporting the idea that cytonemes can act as direct conduits to allow the distribution of a variety of Wnt signalling components (Rogers et al., 2024; Zhang et al., 2024). Membrane scaffolding proteins, such as Flotillin and the Bar protein IRSp53, play major roles in controlling the emergence of these cytonemes (Bischoff et al., 2013; Stanganello et al., 2015; Routledge et al., 2022).

A key unresolved question is how the contacts of these fragile protrusions are stabilised at the receiving membrane. Cytoneme contacts are often transient, suggesting that additional local mechanisms may be required to convert an exploratory membrane encounter into a stable interface. Recently, Ca^2⁺^ transients have been observed at cytoneme contact sites in *Drosophila* that correlate with signalling activity (Huang et al., 2019). In embryonic human stem cells, specialised cytonemes selectively react to Wnts promoting self-renewal, and this contact- dependent response also involves Ca^2⁺^ signalling (Junyent et al., 2020). Similarly, in human cortical neurons, WNT7A-positive dendritic cytonemes induce Ca^2⁺^ signalling, LRP6 clustering and synaptic marker assembly at sites of membrane contact (Piers et al., 2024). Thus, Ca^2⁺^ signalling can accompany effective cytoneme-mediated activation of the Wnt/β-catenin signal; however, its function at the contact sites remains unclear. This raises the possibility that Ca^2⁺^ acts as a local permissive signal enabling efficient Wnt/β-catenin activation at sites of cytoneme contact.

Here, we investigate the role of Ca^2⁺^ signalling at cytoneme contacts and its contribution to canonical Wnt activation in HeLa cells. We show that local Ca^2⁺^ transients at WNT3/3a-positive cytonemes are required to permit and enhance Wnt/β-catenin signalling. Our data support a model in which physical contact between the cytoneme tip and the receiving membrane triggers a local calcium response, stabilises E-cadherin/CDH1 at the contact site and promotes ligand-receptor engagement. These findings identify a contact-dependent step linking physical membrane engagement to chemical pathway activation and provide a mechanism by which receiving cells can discriminate effective Wnt presentation from transient membrane contact and background noise.

## Results

To investigate the mechanism by which Wnt signalling can occur through cytonemes, we used HeLa cells, a human cancer cell line that is well suited for high-resolution imaging-based analysis of Wnt trafficking (Supplementary Fig. 1a). HeLa cells have been used to visualise WNT3a secretion from the secretory pathway to the plasma membrane (Moti et al., 2019). In addition, HeLa cells form abundant filopodia-like surface protrusions, which are known to initiate cell-cell contacts and facilitate signal transmission at the cell cortex (Gonda et al., 1976; Romero et al., 2012; Thompson et al., 2017). We centred the assay on WNT3/3a and the Wnt ligand secretion mediator WLS (Bänziger et al., 2006; Bartscherer et al., 2006) in the Wnt-producing cells, and on FZD10, LRP6 and AXIN1/2 in the receiver cells, because these proteins capture key steps in canonical Wnt signal presentation and are abundantly expressed in HeLa cells (Janda et al., 2012; Willert et al., 2003; Zhan et al., 2017; Nagayama et al., 2002). For high-resolution live imaging with enhanced temporal resolution, we used SIM^2^ microscopy (Piers et al., 2024) to test whether HeLa filopodia carry Wnt pathway components and function as Wnt cytonemes.

### HeLa filopodia carry Wnt ligands, secretion machinery, and receptors

We observed that HeLa cells formed long membrane protrusions that established direct contacts with neighbouring cells. On these filopodia, we found endogenous WNT3 in discrete puncta, and it was strongly enriched at filopodia tips and filopodial contact sites (Fig. 1a; see Supplementary Fig. 1b for antibody control). This localisation was sensitive to Wnt pathway manipulation: WNT3 overexpression increased the number of puncta by 74%, whereas inhibition of Porcupine-mediated Wnt lipidation via Wnt-C59 decreased the proportion of WNT3-positive filopodia by 59% (Fig. 1b, c).

**Figure 1.**
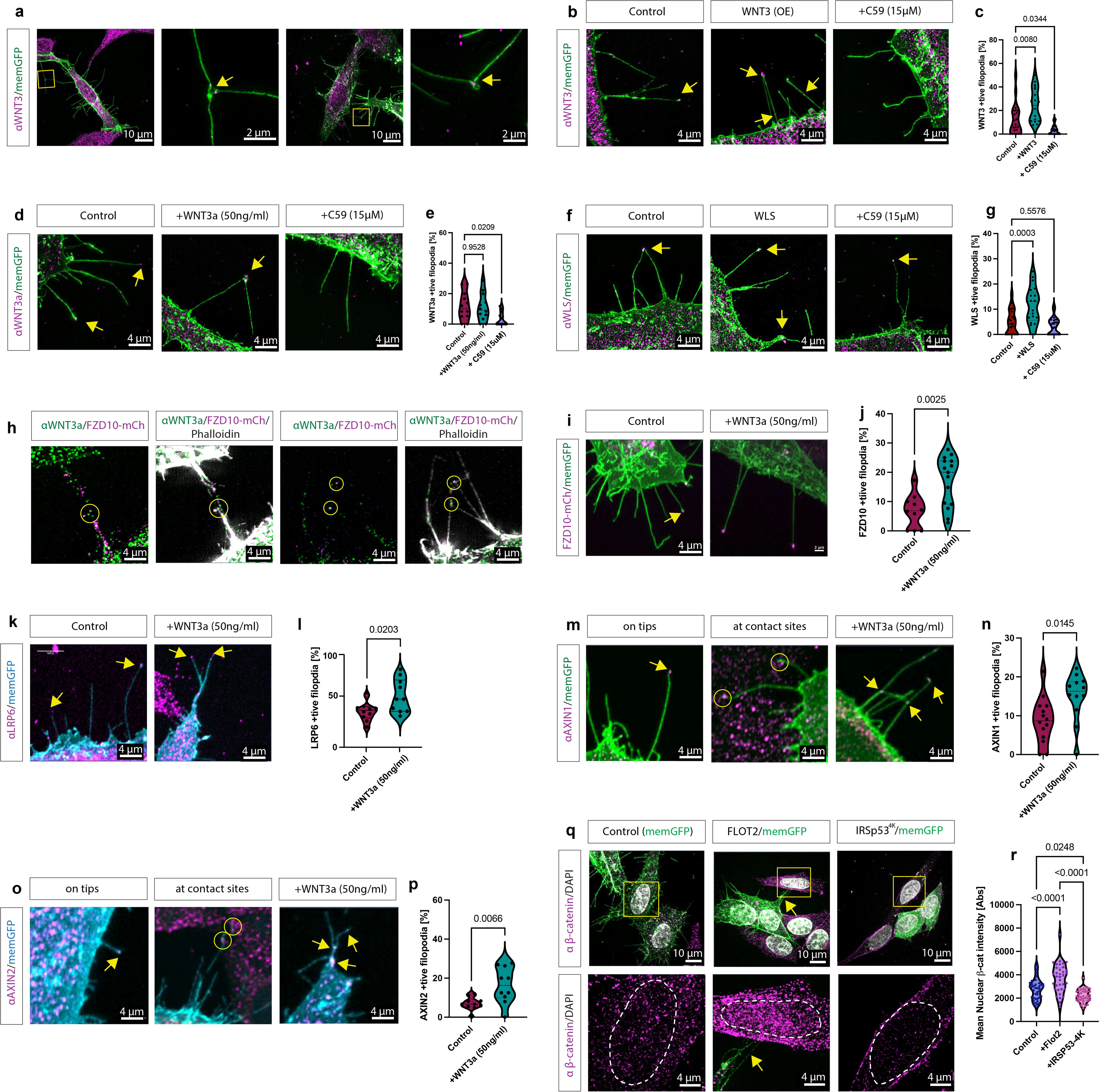
Wnt ligands, receptors and signalosome components accumulate on HeLa cell cytonemes. (a) Immunofluorescence analysis of endogenous WNT3 in membrane-GFP-expressing HeLa cells reveals WNT3 puncta at cytoneme tips and cytoneme-mediated cell–cell contacts. Boxed regions are shown at higher magnification. (b–g) Localisation of endogenous WNT3, WNT3a and WLS on membrane-GFP-positive filopodia. WNT3 and WLS were analysed in control cells, after overexpression of the indicated component, and after treatment with the Porcupine inhibitor Wnt-C59. WNT3a was analysed in control cells, after stimulation with recombinant WNT3a, and after Wnt-C59 treatment. Corresponding quantifications show the percentage of filopodia positive for the indicated Wnt component. (h) Co-localisation of endogenous WNT3a and FZD10-mCherry at phalloidin-positive cytoneme contacts. (i–l) Localisation of the Wnt receptor components FZD10 and LRP6 on membrane-GFP-positive filopodia under control conditions and after recombinant WNT3a stimulation. (m–p) Recruitment of the downstream Wnt signalling components AXIN1 and AXIN2 to membrane-GFP-positive filopodia, cytoneme tips and contact sites, with increased localisation after recombinant WNT3a stimulation. (q,r) Activation of canonical Wnt signalling in HeLa cells expressing the cytoneme regulators FLOT2 or IRSp53^4K^, assessed by nuclear β-catenin staining. Higher-magnification images show the boxed cells; dashed outlines indicate nuclei. All filopodia/cytoneme quantifications show the percentage of membrane-GFP-positive filopodia positive for the indicated protein. Nuclear β-catenin is quantified as mean nuclear β-catenin intensity. Data are presented as violin plots from n = 3 independent experiments, with >10 cells analysed per condition. Statistical significance was determined using Kruskal–Wallis test with Dunn’s multiple comparisons test for WNT3; one-way ANOVA with Dunnett’s multiple comparisons test for WNT3a, WLS and β-catenin; and unpaired two-tailed t-tests for FZD10, LRP6, AXIN1 and AXIN2 comparisons. *p*-values are indicated in the graphs. Yellow arrows indicate positive cytoneme tips or contact sites; yellow circles indicate puncta at contact sites. Scale bars are indicated in each panel.

Next, we tested WNT3a and observed that WNT3a presentation was similarly detectable on these protrusions and decreased by 72% after C59 treatment (Fig. 1d, e; Supplementary Fig. 1c). In contrast, treatment with recombinant WNT3a did not significantly increase filopodia-associated WNT3a levels, indicating that only endogenously generated WNT3a can be transported on these protrusions. In addition to the ligands, we could also detect the endogenous Wnt carrier protein WLS on these filopodia, and WLS overexpression increased the fraction of WLS-positive puncta on protrusions by 89% (Fig. 1f, g, I; Supplementary Fig. 1d) whereas WLS was not reduced upon treatment with Wnt-C59.

Co-labelling experiments showed that WNT3a puncta colocalised with FZD10 at contact sites of these F-actin-positive filopodia (Fig. 1 h), indicating that Wnt ligands and Wnt receptors can be loaded on cytonemes and ligand-receptor complexes at cytoneme-cytoneme contacts as observed previously (Sutton et al., 2026).

As not only the ligand but also the receptors were found on these cell extensions, we next tested whether receptor localisation on cytonemes is Wnt-dependent. When cells were treated with recombinant WNT3a, we significantly enhanced the presentation of FZD10 (Fig. 1i, j) and the co-receptor LRP6 on cytonemes by 153% and 53%, respectively (Fig. 1k, l, Supplementary Fig. 1h), suggesting that Wnt can stabilise and cluster the receptors on filopodia by forming a ligand-receptor complex, most likely the Wnt signalosome (Bilic et al., 2007; Gammons and Bienz, 2018). To test this notion further, we analysed intracellular components of the Wnt signalosome, namely AXIN1 and AXIN2, and observed that these are also localised on tips and on contact sites (Fig. 1m-p, Supplementary Fig. 1f, g, Supplementary Fig. 6i). Furthermore, we found an enrichment of these at contact sites post-treatment with WNT3a protein, with cytoneme localisation of AXIN1 and AXIN2 increasing by 71% and 138%, respectively. Compared with control samples.

We next asked whether filopodia-mediated contacts influence the level of Wnt pathway activation in receiving cells. To this end, we quantified nuclear β- catenin/CTNNB1 accumulation in HeLa cells contacted by neighbouring filopodia (Fig. 1q,r). Cells were seeded at low density to favour discrete filopodia-mediated interactions, and only cells forming visible filopodial contacts were included in the analysis. To modulate contact frequency, we increased filopodia formation by expressing FLOT2 (Routledge et al., 2022) or decreased it by expressing the dominant-negative IRSp53^4K^ construct (Meyen et al., 2015; Stanganello et al., 2015). FLOT2 expression increased the length of filopodia significantly (Supplementary Fig. 1i) and was associated with a significant increase of 34% of nuclear β-catenin levels in the contacted cells (Fig. 1q,r). Conversely, expression of IRSp53^4K^ decreased the number of filopodia (Supplementary Fig. 1i) and led to a significant reduction of 20% in nuclear β-catenin accumulation in contacted cells (Fig. 1q,r).

Based on these results, we conclude that HeLa cells generate specific filopodia that transport Wnt signalling components, which we define as Wnt cytonemes. On the one hand, we can identify Wnt/WLS-positive cytonemes, formed by the Wnt-producing cells, and, on the other, FZD10/LRP6/AXIN1/AXIN2-positive cytonemes, formed by the Wnt-receiving cells. At their contact sites, cytonemes can assemble Wnt signalosomes, which correlate with canonical Wnt pathway activation in receiving cells.

### Calcium transients occur at filopodial contact sites in HeLa cells

Having established that HeLa cells form Wnt cytonemes essential for paracrine signalling, we next asked whether their contact sites exhibit local calcium activity, as has been suggested for other morphogen-carrying cytonemes (Huang et al., 2019). To visualise Ca^2⁺^ dynamics at the plasma membrane, we used the membrane-tethered indicator mem-GCaMP7s (Piers et al., 2024), and we validated its signal behaviour in entire cells and at specific areas of the plasma membrane in HeLa cells over time (Supplementary data 2a-g).

In control cultures, we observed rapid, short-lived, localised Ca^2⁺^ transients at sites of filopodia contact, with signals spatially restricted to the point of interaction (Fig. 2a; Supplementary movie 1). Time-lapse analysis confirmed that these transient Ca^2⁺^ peaks were tightly associated with contact events and were diminished in the absence of cytoneme interaction (Fig. 2b).

**Figure 2.**
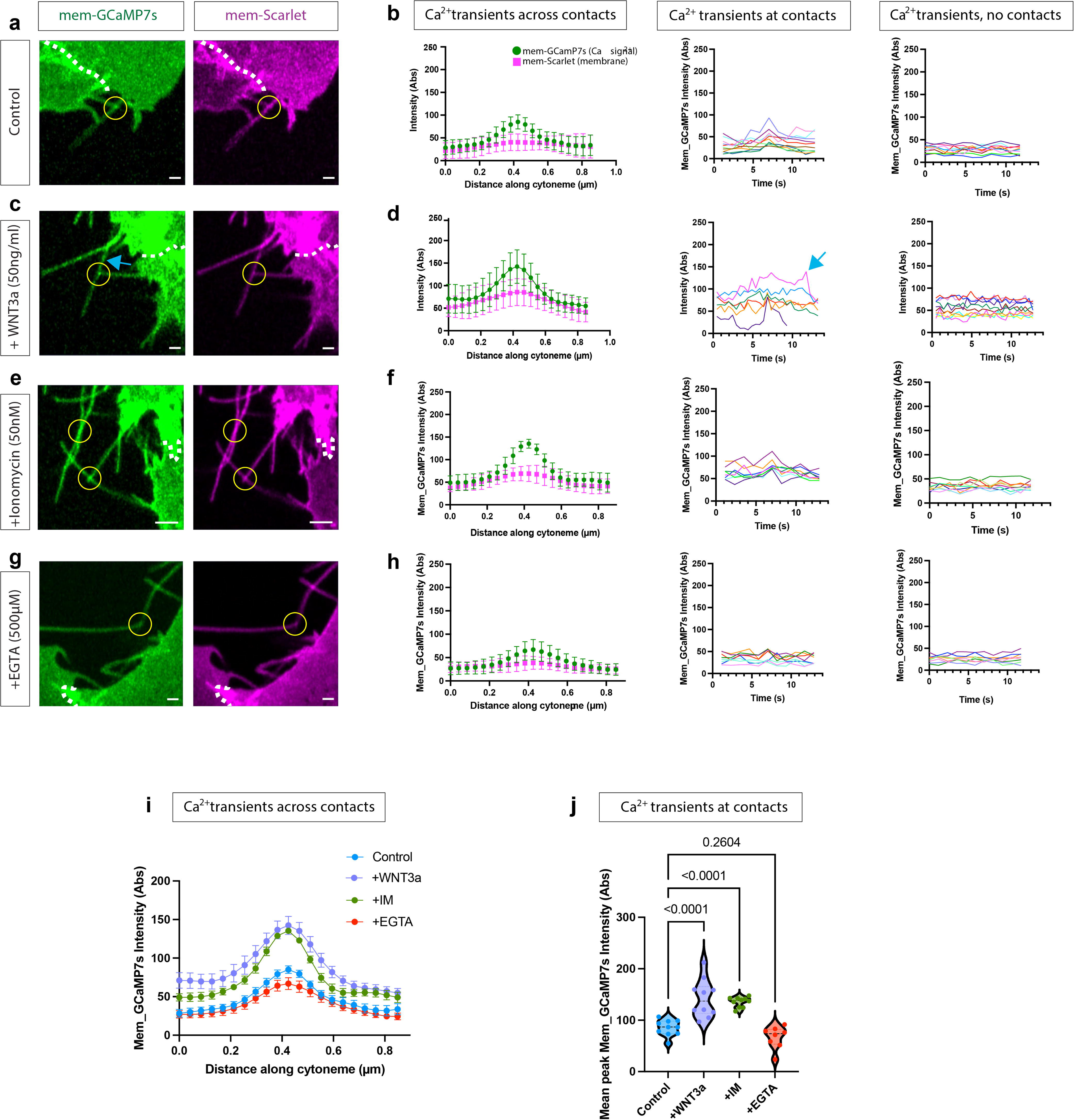
Calcium transients occur at cytoneme-mediated cell–cell contact sites. (a,b) HeLa cells co-expressing membrane-Scarlet and membrane-targeted GCaMP7s were imaged to analyse calcium dynamics at cytoneme-mediated contacts. Representative images show GCaMP7s and membrane signal at contacting cytoneme tips. Quantification shows fluorescence intensity profiles along the distal cytoneme and across the contact site, calcium dynamics over time at contact points, and GCaMP7s signal in non-contacting filopodia. (c–h) Calcium dynamics after modulation of calcium signalling or Wnt pathway activation. Cells were treated with ionomycin (50 nM; c,d), EGTA (500 µM; e,f), or recombinant WNT3a (50 ng/ml; g,h), and analysed as described in (a,b). Line profiles show mean fluorescence intensity along the cytoneme; time-course plots show individual contact events or non-contacting filopodia, as indicated. (i,j) Comparison of contact-associated GCaMP7s signals across conditions. Line profiles show GCaMP7s intensity along cytonemes in control cells and treated cells. Peak GCaMP7s intensity at cytoneme contacts is quantified in (j). Yellow circles mark cytoneme-mediated contact sites; white dashed lines indicate the border between two neighbouring cells. Data are presented as mean ± SEM from n = 3 independent experiments, with >10 cells analysed per condition. Statistical significance in (h) was determined using one-way ANOVA with Dunnett’s multiple comparisons test. p-values are indicated in the graph. Scale bars are indicated in each panel.

We next examined their relationship to Wnt signalling. The addition of recombinant WNT3a enhanced Ca^2⁺^ transients at filopodial contact sites, with a mean peak intensity increase of 68% compared with control samples. (Fig. 2c, d). This indicates that Wnt ligand presentation promotes focal Ca^2⁺^ signalling specifically at their contacts. Interestingly, elevated Ca^2⁺^ levels were also observed along cytonemes in the absence of direct contact, suggesting that WNT3a can stimulate Ca^2⁺^ signalling independently of filopodia interactions (Fig. 2d, blue arrows), potentially through ligand engagement along the filopodia shaft. However, the observed discrete transient peaks remained spatially associated with contact sites, consistent with localised receptor activation.

Finally, we validated whether these transients were Ca^2⁺^-dependent: ionomycin, a Ca^2^⁺ ionophore that elevates cytosolic Ca^2⁺^ by mobilising intracellular stores and promoting Ca^2⁺^ influx across membranes, increased contact-associated mem- GCaMP7s signals (Fig. 2e, f), while EGTA, which chelates extracellular Ca^2⁺^ and lowers free Ca^2⁺^ availability outside the cell, suppressed them (Fig. 2g, h).

Quantification across conditions showed that, similar to WNT3a treatment, Ionomycin treatment strongly elevated Ca^2⁺^ activity at cytoneme contacts by 59%, whereas EGTA treatment produced a significantly lower response than control, with a 21% decrease in mean peak intensity. (Fig. 2i, j). Thus, we conclude that local calcium signalling is a feature of HeLa filopodial contacts and is sensitive to both intracellular and extracellular calcium mobilisation as well as WNT3a stimulation.

### WNT3a and FZD10 increase calcium transients at Wnt cytoneme contacts

To link these contact-associated calcium transients to bona fide Wnt reception sites observed previously (Fig. 1h), we examined the function of the Wnt receptor FZD10 on Ca^2⁺^ transients on HeLa cytonemes. To this end, we overexpressed fluorescently tagged FZD10 and analysed its localisation relative to membrane-associated Ca^2⁺^ signals. FZD10-mCh was readily detected on cytonemes and showed clear enrichment at sites of cytoneme contacts, where it colocalised with focal mem-GCaMP7s transients (Fig. 3a, b). The addition of WNT3a protein further increased both the amplitude and the spatial concentration by two-fold of these contact-associated Ca^2⁺^ signals, and this response remained closely aligned with FZD10-positive contact sites (Fig. 3c, d). Finally, by treating these cells with EGTA to remove extracellular Ca^2⁺^, we could reduce Ca^2⁺^ transients at the FZD10/WNT3a contact sites, suggesting that extracellular Ca^2⁺^ is a regulator of transients at contact sites.

**Figure 3.**
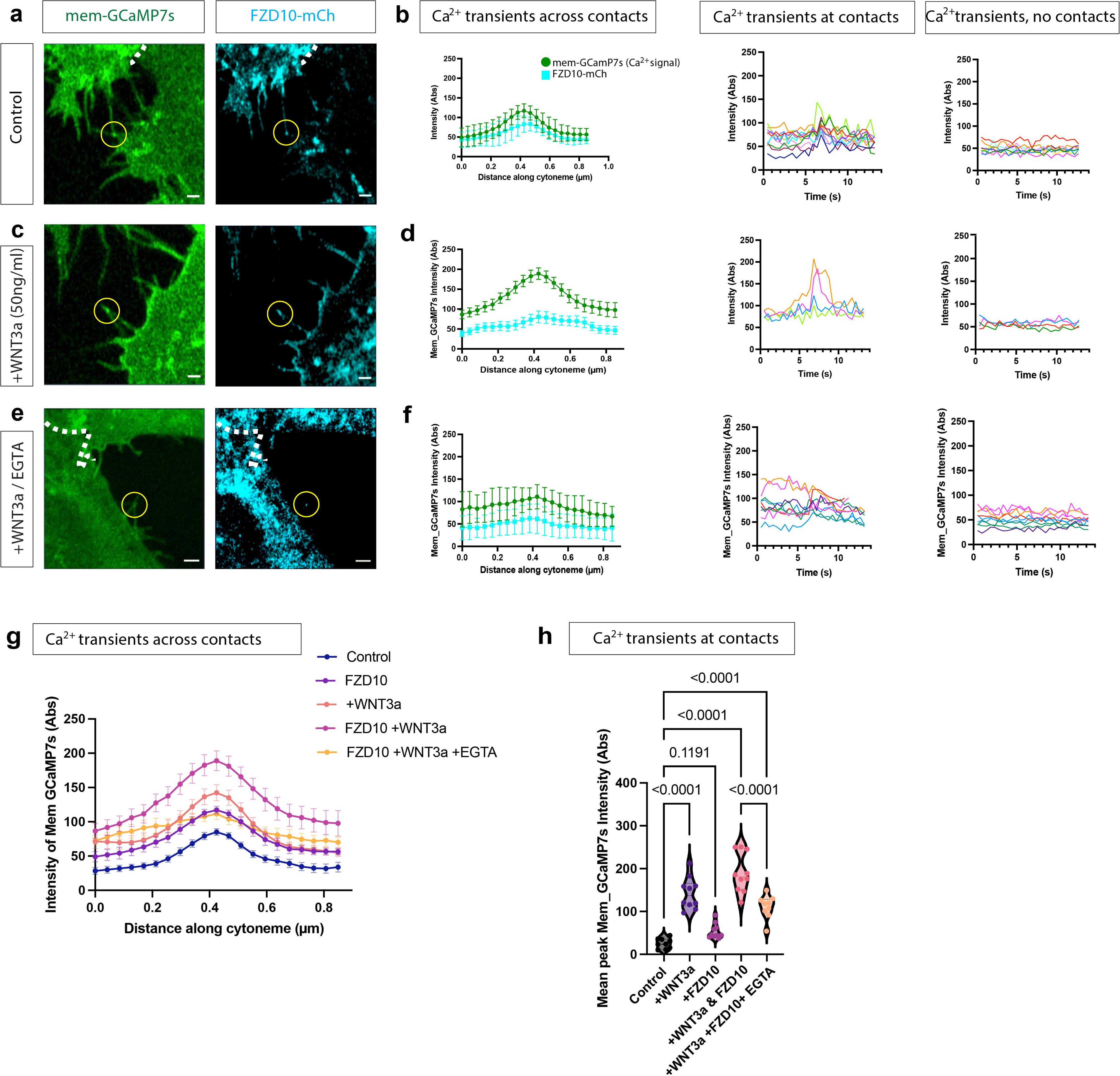
Calcium transient intensity depends on available Wnt pathway components. (a–f) HeLa cells co-expressing FZD10-mCherry and membrane-targeted GCaMP7s were imaged to analyse calcium dynamics at cytoneme-mediated contacts. Cells were analysed under control conditions (a,b), after stimulation with recombinant WNT3a (50 ng/ml; c,d), or after combined WNT3a and EGTA treatment (500 µM; e,f). Quantifications show GCaMP7s and FZD10-mCherry fluorescence intensity profiles along the distal cytoneme and across contact sites, calcium dynamics over time at contact points, and GCaMP7s signal in non-contacting filopodia. (g,h) Comparison of contact-associated GCaMP7s signals across conditions. Line profiles show GCaMP7s intensity along cytonemes in control cells, FZD10-expressing cells, WNT3a- treated cells, FZD10-expressing cells treated with WNT3a, and FZD10-expressing cells treated with WNT3a and EGTA. Peak GCaMP7s intensity at cytoneme contacts is quantified in (h). FZD10 expression and WNT3a stimulation increase local calcium transients, whereas EGTA reduces this response. GCaMP7s and FZD10-mCherry signals are shown as raw fluorescence intensity. Line profiles show mean fluorescence intensity along the cytoneme; time-course plots show individual contact events or non-contacting filopodia, as indicated. Yellow circles mark cytoneme-mediated contact sites white lines indicate the border between two neighbouring cells. Data are presented as mean ± SEM from n = 3 independent experiments, with >10 cells analysed per condition. Statistical significance in (h) was determined using one-way ANOVA with Dunnett’s multiple comparisons test. p-values are indicated in the graph. Scale bars are indicated in each panel.

Quantification across conditions revealed that FZD10 expression did not cause elevated Ca^2⁺^ activity at cytoneme contacts, whereas WNT3a stimulation produced a significantly stronger response, with mean peak intensities increasing by 68%. Importantly, combining FZD10 with WNT3a caused the largest increase of 122% in contact-associated Ca^2⁺^ transients (Fig. 3 g, h). Blocking extracellular calcium with EGTA significantly reduced the WNT3a/FZD10-mediated increase in these transients. Although EGTA treatment can significantly reduce the Ca^2⁺^ transients induced by WNT3a/FZD10, it cannot reduce the transients to the control level, suggesting that the intracellular Ca^2⁺^ pool also contributes to the transients recorded.

Thus, cytonemes form Wnt receptor FZD10-positive contact sites, which are sites of focal calcium activity, and WNT3a and FZD10 act synergistically to enhance Ca^2⁺^ transients at cytoneme contacts. Notably, combined WNT3a treatment and FZD10 overexpression also resulted in reduced cytoneme motility, with protrusions appearing more stable and persistent over time. This observation prompted us to further investigate whether Wnt signalling influences cytoneme contact duration.

### Ca^2⁺^ and Wnt signalling increase cytoneme contact duration

Therefore, we next asked whether Ca^2⁺^ and Wnt signalling regulate the stability of cytoneme contacts between neighbouring HeLa cells. Time-lapse imaging showed that individual cytoneme contacts in control cells were typically short-lived (Fig. 4a, yellow circles) and frequently disengaged (Fig. 4a, yellow arrows) within a 90sec recording period.

**Figure 4.**
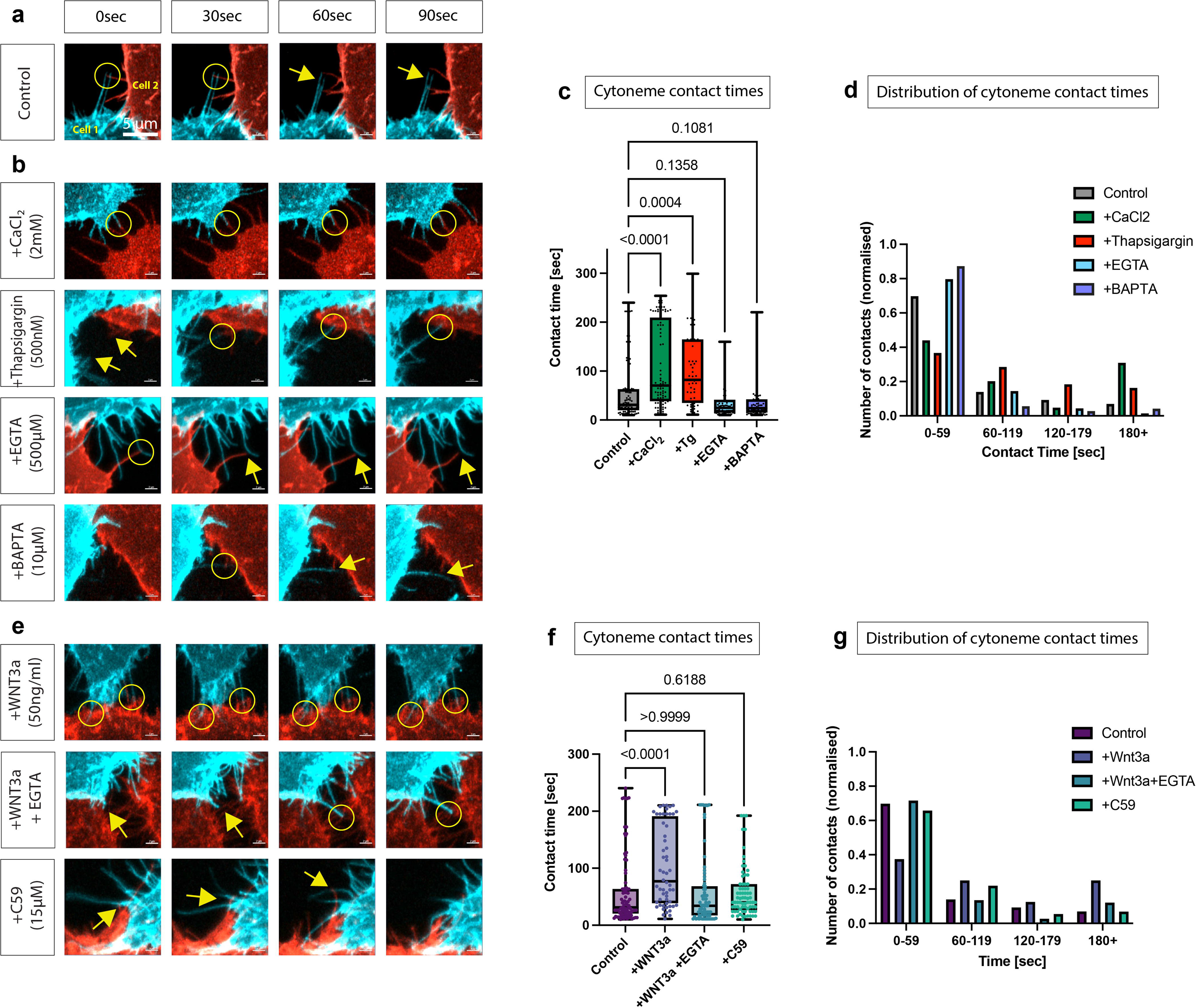
Cytoneme contact duration depends on calcium availability and WNT3a signalling. (a) HeLa cells expressing the indicated membrane markers were co-cultured and live imaged to analyse filopodia-mediated contacts between distinct cell populations. Time-lapse images show the formation, maintenance and disengagement of a representative cytoneme contact. (b–d) Modulation of calcium availability alters contact dynamics. Cells were treated with CaCl (2 mM), thapsigargin (Tg) (500 nM), EGTA (500 µM) or BAPTA-AM (10 µM), and contact duration was quantified from live imaging. Contact times for individual filopodia are shown in (c), and the distribution of contact durations grouped into time intervals is shown in (d). (e–g) WNT3a signalling prolongs cytoneme contact duration. Cells were treated with recombinant WNT3a (50 ng/ml), WNT3a together with EGTA, or the Porcupine inhibitor Wnt- C59 (15 µM). Representative time-lapse images are shown in (e), contact durations are quantified in (f), and contact-time distributions are shown in (g). Cytoneme contact duration was defined as the time from initial contact to disengagement between membrane-labelled cells. Images were acquired every 4.3 s. Data are presented from n = 3 independent experiments, with >10 cells and filopodia analysed per condition. Statistical significance was determined using Kruskal–Wallis test with Dunn’s multiple comparisons test. *p*-values are indicated in the graphs. Yellow circles mark stable cytoneme/filopodia contacts; yellow arrows indicate filopodia tips or contact events. Scale bars are indicated in each panel.

To determine whether calcium availability influences cytoneme contact stability, cells were first treated with pharmacological agents that modulate Ca^2⁺^ levels. First, we found that increasing extracellular Ca^2⁺^ with CaCl significantly prolonged these contacts (84% increase), whereas mobilisation of intracellular Ca^2⁺^ stores with thapsigargin produced a similarly significant increase (72%) in contact persistence (Fig. 4 b-d). Conversely, chelation of extracellular Ca^2⁺^ with EGTA or intracellular Ca^2⁺^ buffering with BAPTA reduced contact duration by 31% and 32%, respectively, and shifted the distribution toward shorter contact events (Fig. 4 b-d). These effects did not reach statistical significance across all conditions, presumably due to the already very short contact times in the control condition.

We next assessed whether Wnt signalling similarly affects cytoneme contact dynamics. We found that the addition of recombinant WNT3a significantly increased cytoneme contact duration compared with control cells (Fig. 4e-g). This WNT3a- induced increase in contact duration was abolished upon co-treatment with EGTA, thereby restoring contact time to levels comparable to those in untreated cells. Similarly, inhibition of the production of active Wnts by Wnt-C59 also reduced contact stability (Fig. 4e-g).

Together, these data show that both Ca^2⁺^ availability and Wnt signalling promote cytoneme contact duration, and the observed Wnt-mediated stabilisation is dependent on Ca^2+^ availability. This data is consistent with a model in which local Wnt activity promotes Ca^2⁺^ signalling, which acts in a positive feedback loop to stabilise the membrane interface for effective signal transfer.

### Ca^2⁺^ and Wnt signalling promote CDH1 accumulation on cytonemes

Having established that cytoneme contact duration is regulated by both Wnt signalling and calcium availability (Fig. 4), we next asked whether the contact- stabilising effects are accompanied by local recruitment of cell adhesion factors to cytonemes. Therefore, we focused our analysis on E-cadherin/CDH1, because it acts as a central, multifunctional signalling hub that links cell-cell adhesion, Wnt signalling, and calcium availability (Heuberger and Birchmeier, 2010). CDH1 also mediates the structural formation of epithelial adherens junctions, maintains tissue integrity, and directly regulates β-catenin availability for both adhesion and Wnt signalling.

Immunofluorescence analysis revealed that endogenous CDH1 can be detected on a subset of membrane-labelled, phalloidin-positive filopodia of HeLa cells and was enriched at their contact sites between neighbouring cells (Fig. 5a; Supplementary Fig. 1e). In addition, overexpression of CDH1-GFP led to a similar accumulation at contact sites (Supplementary Fig. 5). Furthermore, at these contact sites, we also find an accumulation of β-catenin (Fig. 5b), suggesting the formation of adherens junction-like adhesion sites (Nelson and Nusse, 2004), and these sites can be observed at cytoneme-cytoneme contacts and cytoneme-soma contacts (Fig. 5c). In addition, CDH1 was observed to co-localise with endogenous WNT3a at cytoneme tips, suggesting a potential spatial link between adhesion and signalling components at these sites (Fig. 5d).

**Figure 5.**
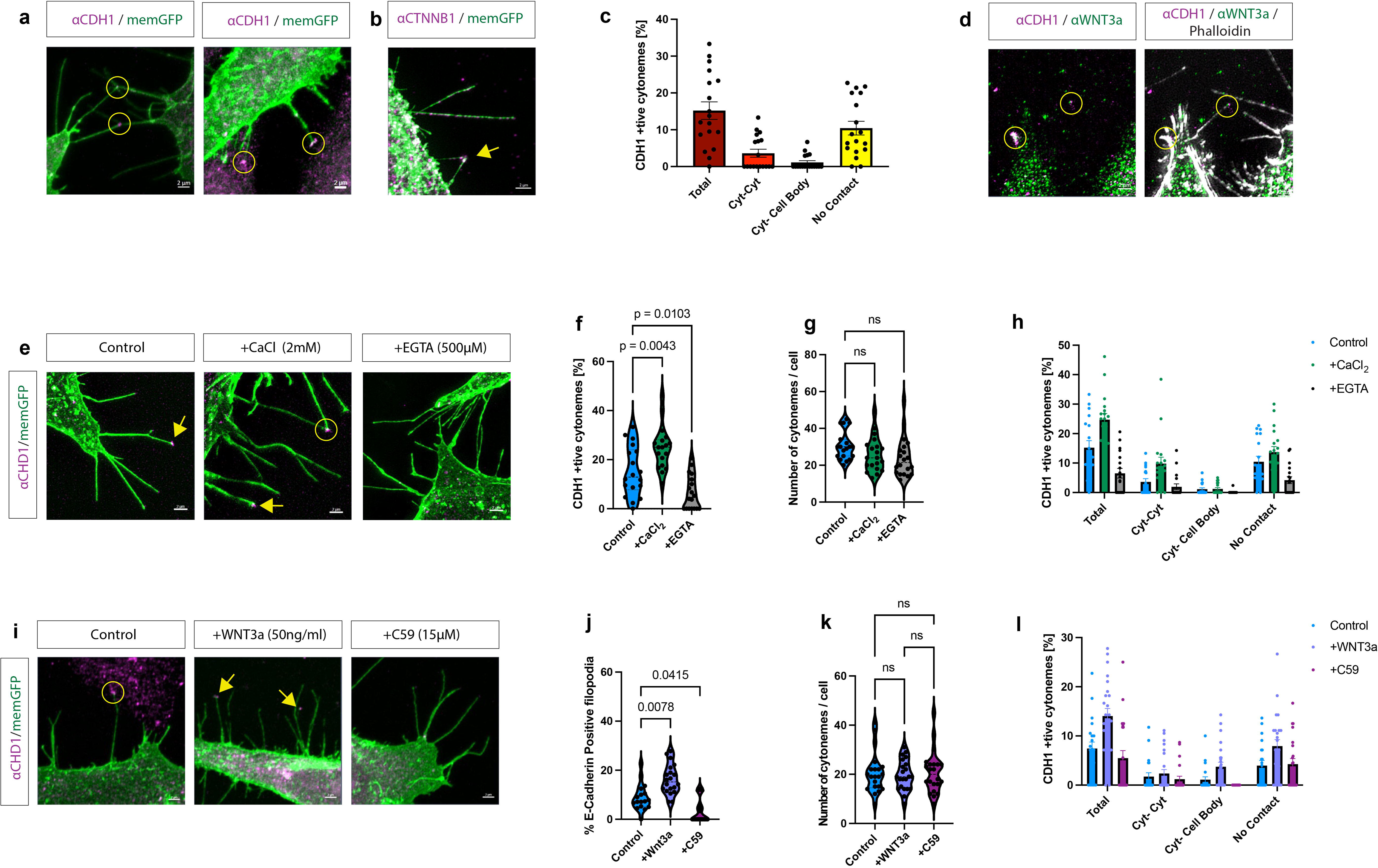
Endogenous CDH1 localises to cytoneme tips and is regulated by calcium and WNT3a signalling. (a–c) Immunofluorescence analysis of endogenous E-cadherin/CDH1 in membrane-GFP-expressing HeLa cells shows CDH1 puncta at cytoneme tips, cytoneme–cytoneme contacts and cytoneme–cell body contacts. β-catenin/CTNNB1 is also detected on membrane-GFP-positive cytonemes. Quantification shows the percentage of CDH1-positive cytonemes in total and according to contact type. (d) Co-localisation of endogenous CDH1 and WNT3a at phalloidin-positive cytoneme contacts. (e–h) Calcium availability regulates CDH1 localisation on cytonemes. Cells were treated with CaCl (2 mM) or EGTA (500 µM), and CDH1 localisation was analysed on membrane-GFP-positive cytonemes. Quantifications show the percentage of CDH1-positive cytonemes, total cytoneme number per cell, and the distribution of CDH1-positive cytonemes according to contact type. (i–l) WNT3a signalling promotes CDH1 localisation on cytonemes. Cells were treated with recombinant WNT3a (50 ng/ml) or the Porcupine inhibitor Wnt-C59 (15 µM). Quantifications show the percentage of CDH1-positive filopodia, total cytoneme number per cell, and the distribution of CDH1-positive cytonemes according to contact type. All panels show HeLa cells under the same imaging conditions unless otherwise indicated. Quantification represents the percentage of CDH1-positive cytonemes/filopodia or the total number of cytonemes per cell, as indicated. Data are presented from n = 3 independent experiments, with >10 cells analysed per condition. Statistical significance was determined using one-way ANOVA with Dunnett’s multiple comparisons test, Brown–Forsythe and Welch ANOVA with Dunnett’s multiple comparisons test, or Kruskal–Wallis test with Dunn’s multiple comparisons test, as appropriate. *p*-values are indicated in the graphs; ns, not significant. Yellow arrows indicate CDH1-positive cytoneme tips or contact sites; yellow circles mark contact-associated puncta. Images were acquired by Zeiss Elyra7 SIM^2^ super-resolution microscopy. Scale bars are indicated in each panel.

We then tested whether Ca^2⁺^ availability affects cytonemal CDH1 localisation. Elevating extracellular Ca^2⁺^ with CaCl□ increased the fraction of CDH1-positive cytonemes by 30%, whereas chelating extracellular Ca^2⁺^ with EGTA reduced this population (Fig. 5e, f) by 50% compared with control samples. By contrast, the total number of cytonemes per cell was not detectably altered under these conditions (Fig. 5g), indicating that Ca^2⁺^ availability may regulate CDH1 recruitment to cytonemes rather than cytoneme abundance. Category-based analysis further showed that CaCl increased, and EGTA reduced, CDH1-positive cytonemes across contact-associated classes (Fig. 5h).

We next asked whether Wnt signalling exerts a similar effect. Recombinant WNT3a increased the proportion of CDH1-positive cytonemes by 95%, whereas inhibition of Wnt secretion with C59 reduced CDH1 localisation to these cytonemes by 24% (Fig. 5i, j). Again, these treatments did not measurably change total cytoneme number per cell (Fig. 5k), but treatment with Wnt protein increased the fraction of CDH1-positive cytonemes within all analysed contact categories (Fig. 5l).

Together, these findings demonstrate that CDH1 is a component of cytoneme contact sites and that its localisation is regulated by both calcium availability and Wnt signalling. This supports a model in which CDH1-mediated adhesion contributes to the stabilisation of cytoneme contacts, potentially linking the presence of Ca^2⁺^ transients to increased Wnt propagation from the producing cell to the receiving cell.

### Ca^2⁺^-dependent cytoneme contacts promote local Wnt pathway activation

We finally asked whether stabilisation of cytoneme-mediated contacts promotes local Wnt signalosome formation and consequently downstream pathway activation. To focus on contact-dependent signalling, HeLa cells were seeded at low density to favour discrete cytoneme-mediated interactions between neighbouring cells, and only cells forming visible cytoneme contacts were included in the analysis, as described above (Fig. 1q).

As a local readout of Wnt pathway activation at the receiving membrane, we quantified the recruitment of the signalosome component AXIN2 to WNT3a-positive cytoneme contact sites. In WNT3a-expressing cells, AXIN2-positive puncta were detected in close proximity to cytoneme-mediated contacts with neighbouring cells, consistent with local signalosome formation at the contact interface (Fig. 6a, Supplementary Fig. 6a,b, i).

**Figure 6.**
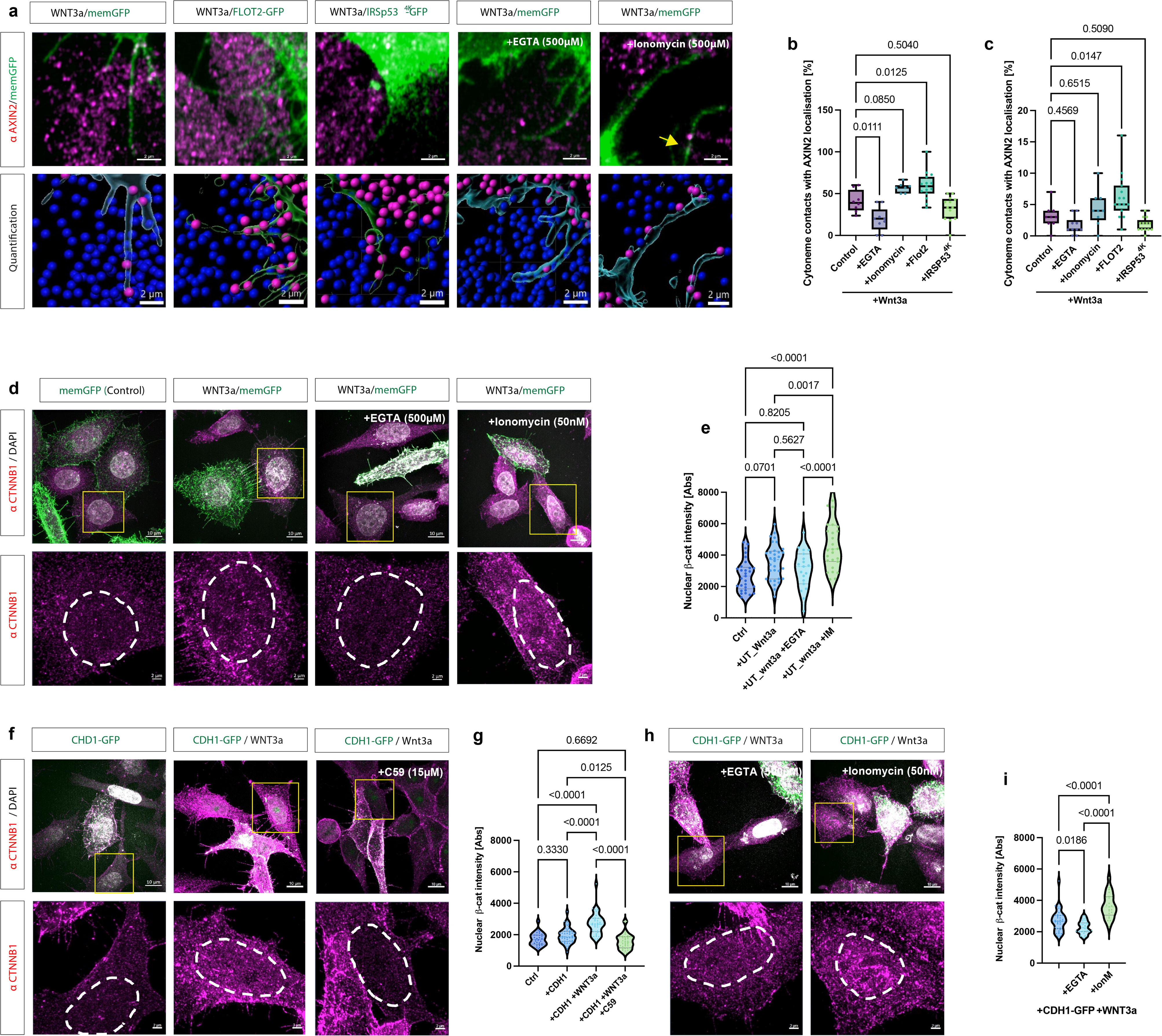
Stabilisation of cytoneme contacts promotes Wnt signalosome formation and canonical Wnt pathway activation. (a–c) Recruitment of AXIN2 to cytoneme contact sites was used as a proxy for local Wnt signalosome formation. Quantification is shown as absolute values (b) and normalised data (b). HeLa cells expressing membrane-GFP were treated with recombinant WNT3a alone or in combination with EGTA, ionomycin, FLOT2- GFP or IRSp53^4K^. Representative images and corresponding segmentation-based quantification show AXIN2-positive puncta at cytoneme contact sites. Quantifications show the percentage of cytoneme contacts with AXIN2 localisation and the number of AXIN2- positive contact sites per cell/field, as indicated. (d,e) Nuclear β-catenin/CTNNB1 localisation was analysed as a downstream readout of canonical Wnt pathway activation. Cells expressing membrane-GFP were analysed under control conditions, after WNT3a stimulation, or after WNT3a stimulation combined with EGTA or ionomycin. Boxed nuclei are shown at higher magnification; dashed outlines surround nuclei. Quantification shows mean nuclear β-catenin intensity. (f,g) Stabilisation of cytoneme contacts by E-cadherin/CDH1 enhances Wnt pathway activation. Cells expressing CDH1-GFP were analysed under control conditions, after WNT3a stimulation, after WNT3a stimulation combined with EGTA, or after treatment with Wnt-C59. Nuclear β-catenin/CTNNB1 was quantified as in (d,e). AXIN2- positive contact sites were identified from segmented membrane-GFP-positive cytonemes and quantified as contact-associated AXIN2 puncta. Nuclear β-catenin was quantified as mean nuclear CTNNB1 fluorescence intensity. Data are presented from n = 3 independent experiments, with >10 cells and cytoneme contacts analysed per condition. Statistical significance was determined using one-way ANOVA with Dunnett’s multiple comparisons test / Kruskal–Wallis test with Dunn’s multiple comparisons test, as appropriate. *p*-values are indicated in the graphs. Yellow arrows indicate AXIN2-positive cytoneme contact sites; yellow boxes indicate regions shown at higher magnification; dashed outlines mark nuclei. Scale bars are indicated in each panel.

Increasing cytoneme formation by FLOT2 expression enhanced the number of cytonemes with AXIN2-positive signalosome formation at contacts, whereas reducing cytoneme formation by expression of IRSp53^4K^ decreased this response (Fig. 6b, c). Although treatment of HeLa cells with recombinant WNT3a or the AXIN stabiliser IWR1 increased AXIN2, only WNT3a treatment could increase AXIN2 on cytonemes (Supplementary Fig. 6a-e), suggesting that externally provided WNT3a can recruit AXIN2 to cytonemes and presumably induce a receiver cytoneme. Next, we asked if calcium availability impacts cytoneme-mediated signalosome induction. We observed that chelating extracellular Ca^2⁺^ with EGTA significantly reduced the fraction of cytoneme contacts associated with AXIN2 localisation by 33%, whereas stimulation of internal Ca^2⁺^ signalling with ionomycin significantly increased AXIN2 recruitment to contact sites by 36% (Fig. 6a-c). These results indicate that local Wnt signalosome assembly at cytoneme contacts is sensitive to Ca^2⁺^ availability.

We then asked whether these local contact-associated signalling events translate into downstream activation of the canonical Wnt pathway. Nuclear β- catenin/CTNNB1 accumulation was quantified in cells targeted by cytonemes. Compared with membrane-GFP control cells, WNT3a expression increased nuclear CTNNB1 levels in cytoneme-targeted neighbour cells by 31% (Fig. 6d,e). Similar to cells exposed to recombinant WNT3a (Supplementary Fig. 6f,g). The increased CTNNB1 accumulation could be reduced by EGTA treatment, indicating that extracellular Ca^2⁺^ is required for efficient cytoneme-mediated Wnt signalling. Conversely, ionomycin treatment further increased nuclear CTNNB1 accumulation by 79% compared to controls, consistent with a positive effect of Ca^2⁺^ signalling on canonical Wnt pathway activation downstream of cytoneme contacts. We confirmed these findings by using recombinant WNT3a (Supplementary Fig. 6h, i).

Finally, we tested whether CDH1-mediated stabilisation of cytoneme contacts could provide a mechanistic explanation for enhanced signalling. Co-overexpression of CDH1-GFP and WNT3a significantly increased nuclear CTNNB1 accumulation in neighbouring cytoneme-targeted cells by 57% compared with control (Fig. 6f,g). This increase was significantly reduced by treatment with the Porcupine inhibitor C59, indicating that CDH1-dependent contact stabilisation acts together with Wnt presentation to enhance pathway activation in receiving cells. We obtained similar results upon stimulation with recombinant WNT3a (Supplementary Fig. 6j,k).

We then tested whether this effect depends on calcium availability. We find that chelating of extracellular Ca^2⁺^ with EGTA significantly reduced nuclear CTNNB1 accumulation by 16%, whereas treatment with ionomycin significantly enhanced it by 35% in cytoneme-targeted cells (Fig. 6h,i).

Together, these data support a model in which Ca^2⁺^-dependent stabilisation of cytoneme contacts promotes Wnt signal transfer. Increased Ca^2⁺^ availability, WNT3a stimulation, and CDH1-mediated contact stabilisation enhance AXIN2-positive signalosome formation at cytoneme contact sites and increase nuclear CTNNB1 accumulation in contacted cells, whereas reducing Ca^2⁺^ availability or Wnt secretion suppresses these responses. We conclude that Calcium/CDH1 acts as a gatekeeper for cytoneme-mediated Wnt signalling. Although this analysis focuses on cells connected by visible cytoneme-mediated contacts and thus supports a contact-dependent mechanism, it does not rule out additional contributions from other paracrine modes of Wnt exchange.

## Discussion

In this work, we show that HeLa filopodia carry core Wnt pathway components and can function as Wnt cytonemes. Wnt cytoneme contacts exhibit focal Ca^2⁺^ transients, and WNT3a and FZD10 enhance this local calcium activity. Mechanistically, we find that Ca^2⁺^ and Wnt signalling prolong cytoneme contact duration by promoting local CDH1-mediated adhesion of cytonemes and increasing local and downstream canonical Wnt pathway activation in neighbouring cells. These data support a model in which local Ca^2⁺^ signalling works synergistically with CDH1 to stabilise cytoneme contacts and thereby facilitate prolonged Wnt ligand transfer and efficient Wnt/β-catenin signalling.

### Are the Wnt/Ca^2⁺^ and Wnt/β-catenin pathways always antagonistic?

The Wnt network consists of various signalling branches that are thought to function in a mutually repressive manner. In particular, the Wnt/β-catenin and the Wnt/Ca^2⁺^ pathways are believed to act antagonistically: Wnt5a can suppress β- catenin-dependent signalling by promoting β-catenin degradation, and it can inhibit Wnt3a-driven TCF/LEF transcriptional responses under defined receptor conditions (Mikels and Nusse, 2006; Nemeth et al., 2007). However, Wnt5a can also support β- catenin-dependent transcription in a different receptor context (Mikels and Nusse, 2006; van Amerongen et al., 2012). Complementary, β-catenin-dependent Wnt ligands can induce intracellular calcium responses, as shown for Wnt3a in hippocampal neurons (Avila et al., 2010). Furthermore, during zebrafish heart morphogenesis, Wnt9a/b-dependent tissue remodelling requires combined canonical and non-canonical pathway activity by the very same ligands, rather than strict engagement of only one branch (Paolini et al., 2023). Together, these findings indicate that Wnt/β-catenin and Wnt/Ca^2⁺^ signalling can act antagonistically but also cooperatively, depending on developmental and cellular context, as well as on different spatial and temporal scales.

### Is the function of the cytoneme synapse dependent on Ca^2⁺^ signalling?

Local Ca^2⁺^ transients are a recurrent feature of cytoneme-mediated signalling. In the *Drosophila* air sac primordium, dpp/BMP cytoneme contacts use synapse-like glutamatergic signalling in which glutamate released from signalling cells activates AMPA-type receptors on cytonemes and triggers transient Ca^2⁺^ elevations required for signal uptake and target-tissue development (Huang et al., 2019). Cytonemes can also propagate local calcium-firing activity over long distances through embryonic tissues to regulate BMP gradient formation and embryonic polarity in chick embryos (Lee et al., 2024). These studies suggest that cytoneme contacts are functional morphogen signalling interfaces rather than passive membrane appositions (Kornberg and Roy, 2014). A similar principle has been observed in mammalian stem cells. Embryonic stem cell cytonemes selectively engage localised self-renewal Wnt sources, and this response depends on Wnt receptors, ionotropic glutamate receptors, and localised Ca^2⁺^ transients (Junyent et al., 2020). These Ca^2⁺^ signals are involved in stem cell-niche pairing, which can lead to Wnt/β-catenin pathway activation. Finally, this logic extends to neuronal protrusions. We have previously shown that WNT7a-bearing dendritic cytonemes in human cortical neurons accumulate WNT7a at contact sites and show focal Ca^2⁺^ transients together with LRP6 clustering, synaptic marker assembly, and spine maturation (Piers et al., 2024). Thus, local Ca^2⁺^ signalling is linked to spatially restricted morphogen presentation by cytonemes in multiple systems.

Our findings place local Ca^2⁺^ signalling upstream of Wnt/β-catenin activation at cytoneme contact sites and, therefore, add important spatial regulation to the Wnt signalling outputs. This does not conflict with earlier studies showing antagonism between Wnt/β-catenin and Wnt/Ca^2⁺^ signalling at the level of whole cells, tissues or developmental systems. Rather, it indicates that these pathway relationships are organised across different temporal scales. System-level antagonism is typically inferred from integrated outputs, such as changes in β-catenin stability, transcriptional responses, or tissue patterning phenotypes, measured over extended periods. By contrast, our analysis resolves the earliest events at the membrane interface between signal-producing and receiving cells. At this scale, Ca^2⁺^ does not act as a global inhibitor of canonical Wnt signalling, but as a local permissive signal that appears to convert transient cytoneme contact into a signalling-competent state. We propose that the Ca^2⁺^ transient triggered upon cytoneme contact promotes the assembly of a ligand-receptor complex at the cytoneme contact sites. This would create a local membrane environment in which a Wnt signalosome can form and operate efficiently and repeatedly.

Our data imply that Wnt reception is not determined solely by ligand availability or receptor expression, but by a contact-dependent gating step at the cytoneme synapse. Second, it explains why conclusions drawn from whole-cell or tissue-level readouts cannot be transferred directly to the subcellular level. A Ca^2⁺^ signal that facilitates canonical Wnt signalling activation within a restricted membrane microdomain may still contribute to antagonistic pathway behaviour when integrated across the entire cell over longer time scales.

### How are Ca^2+^ transients at the cytoneme tips mechanistically linked to the activation of the Wnt/β-catenin signalling cascade?

Classical cadherins form adhesive contacts between neighbouring cells, generating stable membrane interfaces that support local signalling and the transfer of membrane-associated components across the contact site (Gumbiner, 2005; Nelson et al., 2013). For example, E-cadherin/CDH1 is a Ca^2⁺^-dependent adhesion receptor that nucleates adherens junctions and couples cell-cell contacts to the cortical actin cytoskeleton through catenins (Niessen et al., 2011). This places CDH1 in a prime position to influence local Wnt reception, as β-catenin functions both as a junctional component downstream of CDH1 and as the central effector of canonical Wnt signalling (Nelson and Nusse, 2004; Valenta et al., 2012). Wnt inputs can modulate this adhesive interface in multiple ways. Canonical Wnt signalling changes the availability and distribution of β-catenin, whereas non-canonical Wnt ligands such as Wnt5a have been reported to promote Ca^2⁺^-dependent cell adhesion and to enhance CDH1-β-catenin complex formation (Heuberger and Birchmeier, 2010; Medrek et al., 2009). In parallel, Ca^2⁺^ acts directly on CDH1 at two levels: extracellular Ca^2⁺^ is required for cadherin trans-interaction, and local intracellular Ca^2⁺^ signals regulate contact assembly and remodelling through calcium-sensitive control of junctional and actin-associated components (Gumbiner, 2005; Kim et al., 2011). Our data places these activities into a common local framework. We propose that cytoneme-delivered Wnt triggers a focal Ca^2⁺^ signal at the receiving membrane, which stabilises CDH1 across the contact site and thereby promotes productive ligand–receptor engagement. In support of the dual function of Ca^2⁺^ on CDH1, we found that modulating extracellular Ca^2+^ (through EGTA and CaCl_2_) and cytosolic Ca^2+^ levels (through thapsigargin, ionomycin, and BAPTA) have similar effects on cytoneme contact stability.

Based on the observed results, we suggest that Wnt provides the instructive signal, Ca^2⁺^ provides the local enabling input, and CDH1 provides the stabilising adhesive scaffold. Thus, at cytoneme contacts, Wnt, Ca^2⁺^ and CDH1 act in a coordinated and mutually reinforcing manner. Our model is supported by emerging evidence that CDH1 has a role in adhesive stabilisation at signalling contacts. In mouse stem cells, transient CDH1 overexpression improved cytoneme-dependent pairing with trophoblast stem cells (Junyent et al., 2021), whereas in the *Drosophila* muscle progenitor niche, “niche-adhering” cytonemes require receptor-dependent adhesion to maintain niche occupancy and to receive contact-dependent signals (Patel et al., 2022).

In summary, we suggest that CDH1 is not simply part of a generic adhesive scaffold, but part of the mechanism by which the receiving membrane authenticates Wnt ligand-carrying cytoneme contact. Only contacts that trigger the local Ca^2⁺^ response would be stabilised sufficiently to support robust ligand presentation, receptor clustering and downstream β-catenin activation. Contacts lacking this local Ca^2⁺^-dependent reinforcement would remain transient and fail to mature into signalling hubs.

### Ca^2⁺^ signalling in cell-connecting protrusions

The role of Calcium has also been studied in other cellular extensions. For example, tunnelling nanotubes (TNTs). TNTs are generally described as open-ended membranous bridges that directly connect the cytoplasm of two cells and support the transfer of ions, small molecules, vesicles, organelles, and electrical activity over distance (Gerdes and Carvalho, 2008; Rustom et al., 2004; Wang et al., 2010). Structural work has reinforced this view by showing that TNTs can contain continuous membrane channels and cytoskeletal elements compatible with direct intercellular coupling, rather than transient apposition alone (Cordero Cervantes and Zurzolo, 2021; Sartori-Rupp et al., 2019). Within this framework, Ca^2⁺^ signalling has been linked to propagated intercellular activity and to IP receptor-dependent amplification within the nanotube, consistent with a mode of communication in which two cells become transiently integrated into a shared conductive unit (Austefjord et al., 2014; Smith et al., 2011; Wang and Gerdes, 2012). Work in neurons further showed that the Wnt/Ca^2⁺^ pathway promotes TNT formation and TNT-mediated cargo transfer, with Wnt5a acting upstream of CaMKII-dependent cytoskeletal regulation (Vargas et al., 2019). Together, these studies place TNT-associated Ca^2⁺^ signalling in the context of long-range coupling, cargo exchange, and pathway-dependent stabilisation of persistent intercellular bridges.

Our data support a different role for Ca^2⁺^ at cytoneme contacts. Here, Ca^2⁺^ signals are focal, short-lived, and restricted to the membrane interface at which Wnt presentation occurs. They do not resemble propagated intercellular currents, but instead mark a local event at the site of ligand reception. We, therefore, favour a model in which Ca^2⁺^ acts as a contact-dependent gating signal in cytoneme-mediated signalling.

## Conclusion

In conclusion, our data support the view that cytoneme-mediated signalling is regulated by a contact-authentication mechanism. In this framework, physical membrane engagement, local Ca^2⁺^ signalling, and CDH1 stabilisation form a coupled module that determines whether a cytoneme contact is simply exploratory or becomes a signalling synapse (Huang et al., 2019). At this signalling synapse, focal Ca^2⁺^ signalling acts as a contact-authentication mechanism, and CDH1-mediated adhesion can accompany Wnt reception and, consequently, determine how efficiently ligand handover can occur across the membrane interface. Thus, local adhesion and calcium signalling may set the gain of cytoneme-mediated Wnt signalling by controlling the efficiency of ligand delivery at individual contact sites.

## Acknowledgement

Research in the Scholpp lab is supported by the BBSRC (Research Grants BB/S016295/1 and OPP490 as well as a BBSRC Equipment grant, BB/T017899/1), by the Wellcome Trust Discovery Award 8438235 and by the Living Systems Institute, University of Exeter. E.C. is supported by a predoctoral fellowship from the SWBio GW4 PhD network. We would further like to thank Gary Davidson (KIT), Cerys Manning (University of Manchester) and the entire Scholpp lab for their critical comments on the manuscript and the Exeter Bioimaging Centre for their support.

## Material and Methods

### Cell culture maintenance

The HeLa cell line (kindly provided by the Costello lab, Exeter) was maintained in antibiotic-free high-glucose DMEM (ThermoFisher Scientific) supplemented with 10% fetal bovine serum (FBS). Cells were cultured at 37 °C in a humidified atmosphere containing 5% CO and were routinely passaged at approximately 80% confluency using 0.01% trypsin (ThermoFisher Scientific). For fixed imaging experiments, cells were seeded onto #1.5 glass coverslips at a density of 2–4 × 10□ cells per well. Cells were routinely tested for mycoplasma contamination using endpoint PCR testing every three months and broth-based testing annually

### Cell culture transfection

Cells were transiently transfected using a reverse transfection approach with FuGENE HD (Promega) at a reagent-to-DNA ratio of 3:1, with a final DNA concentration from 50–1000 ng/mL. For co-culture experiments, cell populations were transfected independently as described above. After 24 hours, cells were trypsinised, counted using a haemocytometer, and reseeded together at the indicated ratios. Cells were imaged 24 hours after mixing. The following DNA plasmids were used pCAG-mGFP membrane bound GFP (Addgene #43816), PCS2+-Frizzled10-mCherry [zFrizzled10 cDNA cloned into PCS2+-Wnt8a-mCh using ECoRI and BAMHI], pCS2+-GAP43-jGCaMP7s [cloned from pGP-CMV- jGCaMP7s (Addgene #104463) into pCS2+-GAP43-GFP using XBal/SnaBl], PCS2+- LCK-mScarlet3 [Cloned from pLCK-mScarlet3_C1 (Addgene #189771) into PCS2+ vector using Xbal and Clal], pcDNA-Wnt3 (Addgene #35909), pcDNA- Wnt3a (Addgene #35908), PCS2+-WLS [cloned from pcDNA-WLS-mEGFP (*M.Boutros*, DKFZ, Heidelberg, Germany) into PCS2+ vector using Cla1 and EcoR1), hE- cadherin-pcDNA3 (Addgene #45769), E-Cadherin-GFP (Addgene #28009).

### Antibody Staining

Cells were seeded onto 1.5 square coverslips for 24 hours with pharmacological treatment if necessary. Cells were washed with 1xPBS and fixed with modified Mem-Fix (4% formaldehyede, 0.2% glutaraldehyde, Sorenson’s phosphate buffer, pH7.4) (Bodeen et al., 2017; Rogers and Scholpp, 2021) (7 minutes, 4°C). The cells were the permeabilised in a blocking buffer at room temperature for 45 minutes. Coverslips were incubated in primary antibody diluted in incubation buffer overnight at 4°C. The following primary antibodies were used: Rabbit anti-Wnt3 (1:100, Abcam #ab32249), Mouse anti-Wnt3a (1:100, Abcam #ab169175), Anti Lrp6 (1:100, Cell Signaling Tech. #3395S), rabbit anti-Axin1 (1:200, ProteinTech #16541-1-AP), rabbit anti-Axin2 (1:200, ProteinTech #20540-1-AP), anti-CD324 (1:100, Invitrogen #14- 3249-82), mouse anti-GPR177 (1:50, Merck #MAB587), rabbit β-catenin (1:200, ProteinTech #51067-2-AP). After washing in PBS, cells were incubated in secondary antibody at room temperature for 1 hour in the dark. The following secondary antibodies were used: Goat anti-rabbit 568 (1:1000, Abcam), donkey anti-mouse (1:1000, Abcam), donkey anti-rat 568 (1:1000, Abcam). Nuclei were counterstained with DAPI (300nM), and F-actin were counterstained with Alexafluor phalloidin 647 (1:1000, Thermofisher #A30107). Coverslips were mounted onto Glass slides using ProLong diamond anti-fade mountant (Invitrogen) and incubated at 4°C overnight. Samples were imaged as described below.

### DNA preparation

Plasmids were amplified in *Escherichia coli* using standard transformation procedures. Briefly, 2 µL of plasmid DNA was transformed into 20 µL of chemically competent *E. coli*, incubated on ice for 20 minutes, heat-shocked at 42 °C for 30 seconds, and returned to ice for 2 minutes. SOC medium was added, and cells were recovered at 37 °C with shaking at 300 rpm for 1 hour. Transformed bacteria were plated onto selective agar plates containing either ampicillin (100µg/mL) or kanamycin (5µg/mL) and incubated overnight at 37 °C. Individual colonies were expanded in LB medium supplemented with the appropriate antibiotic at 37 °C with shaking at 300 rpm for 24 hours. Plasmid DNA was isolated using a MaxiPrep kit (Qiagen) according to the manufacturer’s instructions. Purified DNA was eluted in 100 µL endotoxin-free buffer. DNA concentration and purity were determined using a NanoDrop spectrophotometer. Endo-toxin free DNA was used for all transfection experiments.

### Pharmacological drug treatment

Pharmacological agents were used to manipulate Wnt signalling and intracellular calcium dynamics. Wnt signalling was inhibited using IWR-1 (2–10 µM), or Wnt-C59 (100–500 nM). Calcium availability was manipulated using multiple approaches. Extracellular calcium was reduced using EGTA (2–5 mM). Both extracellular and intracellular calcium were chelated using BAPTA-AM (5–20 µM). Intracellular calcium release from endoplasmic reticulum stores was induced using thapsigargin (1 µM), and extracellular calcium influx was stimulated using ionomycin (1–5 µM). Cells were treated with the indicated agents for 4–24 hours prior to live-cell imaging or fixation, as specified for each experiment. Equivalent vehicle controls were included where appropriate.

### Imaging techniques

Fixed samples were mounted onto glass slides prior and were imaged using either Imaging was performed using a Zeiss Elyra7 with SIM^2^ super-resolution technology or a Zeiss Airyscan 880 microscope. For Airyscan imaging of fixed samples, images were acquired using a 63× oil immersion objective (NA 1.6) in Airyscan Fast mode. Raw data was processed using the Airyscan reconstruction algorithm in Zeiss Zen software and z-stacks were rendered as maximum intensity projections (MIPs). Structured illumination microscopy (SIM) was performed on the Zeiss Elyra 7 SIM^2^ system using a 63× oil immersion objective (NA 1.64). Image stacks were reconstructed using either SIM or SIM^2^ processing in Zeiss Zen software prior to the generation of maximum intensity projections. Live-cell imaging was performed on the Airyscan microscope using a 63× oil immersion objective (NA 1.6). Cells were maintained at 37 °C and 5% CO throughout. Time-lapse z-stacks were acquired in Airyscan Fast mode. For calcium imaging, 2-4 z-planes were acquired every 1.17 seconds. Across all experiments, excitation was performed using 405, 488, 561, or 647 nm laser lines as appropriate. image reconstruction and processing were applied uniformly across experimental conditions.

### Statistical analysis

Quantitative image analysis was performed using Fiji (ImageJ) and Imaris software. Fiji was used for quantification of puncta localisation and distribution along cytonemes. For puncta analysis, a spots surface was created within Imaris and filtered using an intensity threshold calculated from the mean background fluorescent intensity plus two standard deviations (mean +2SD).Only spots that exceeded this threshold were included within subsequent analyses. Membrane localisation was quantified using Imaris through three-dimensional surface reconstruction. Statistical analyses were performed using GraphPad Prism. Data normality was assessed prior to testing. Parametric data were analysed using unpaired two-tailed Student’s *t*-tests or one-way analysis of variance (ANOVA) with Dunnett’s post-hoc test for multiple comparisons. Non-parametric data were analysed using Kruskal–Wallis tests with Dunn’s post-hoc correction. Unless otherwise stated, *n* represents the number of independently analysed cells from at least three biological replicates. All tests were two-sided, and a *p* value of <0.05 was considered statistically significant.

## Notes

### Competing Interest Statement

The authors have declared no competing interest.

