## Supplementary Data for "Calcium signals at cytoneme contacts amplify Wnt/β- catenin signalling"

**Supplementary Figures**

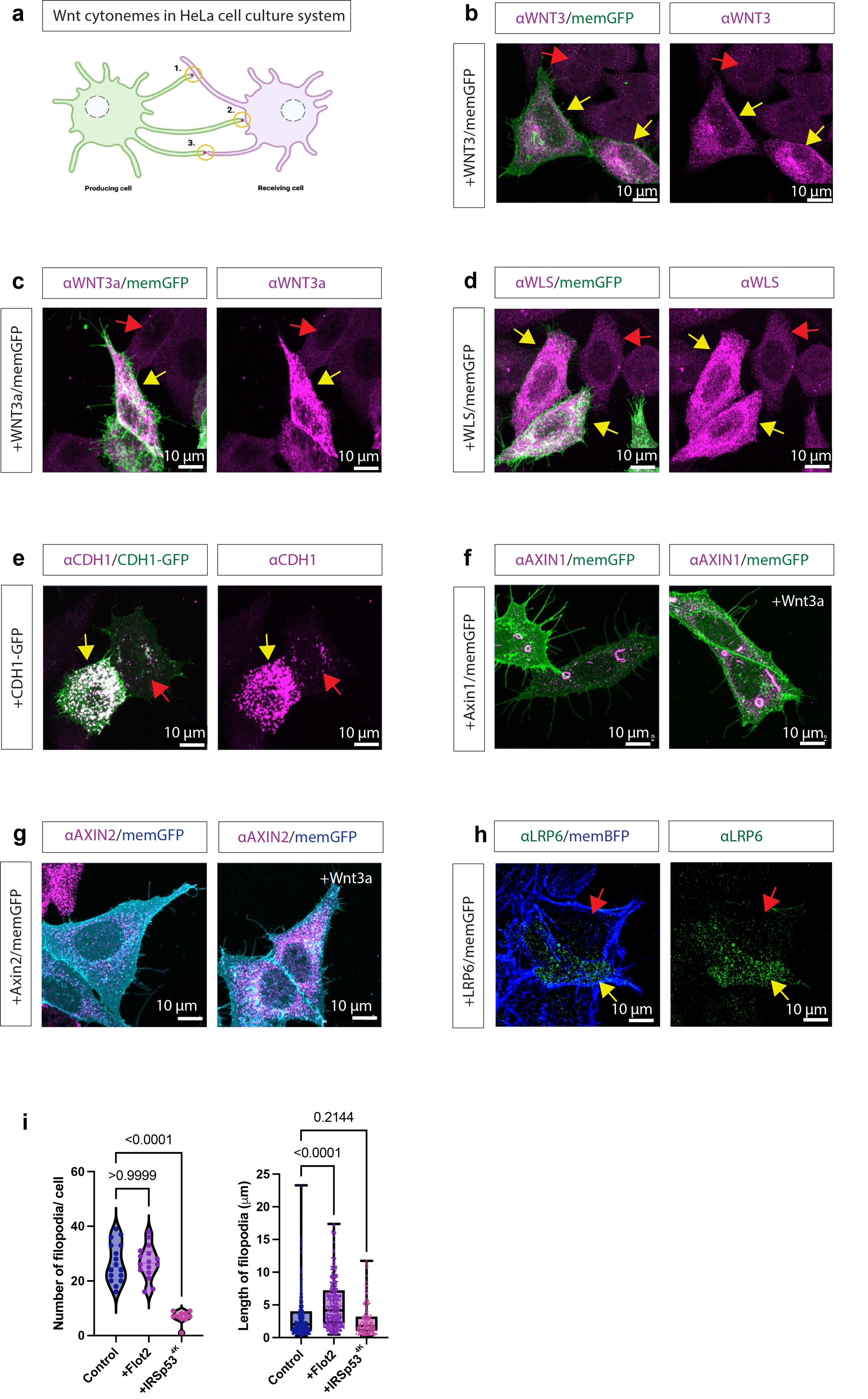

**Supplementary Figure 1. Validation of antibodies of Wnt pathway components in the HeLa cytoneme culture system.** (a) Schematic of the HeLa co-culture system used to analyse Wnt-associated cytoneme contacts between producing and receiving cells. Cytoneme contacts were classified as cytoneme–cytoneme contacts, cytoneme–cell body contacts, or non-contacting cytonemes. (b–e) Immunofluorescence control analysis of WNT pathway components in membrane-GFP-expressing HeLa cells. Yellow arrows indicate cells co-transfected with the same Wnt component used in the immunofluorescence experiment, whereas red arrows indicate non-transfected control cells. (f,g) Endogenous AXIN1 and AXIN2 localisation in HeLa cells under control conditions and after recombinant WNT3a stimulation. WNT3a promotes recruitment of AXIN1 and AXIN2 to membrane-associated puncta.
(h) LRP6 antibody staining in membrane-BFP-expressing HeLa cells. Yellow arrows indicate cells co-transfected with the same Wnt component used in the immunofluorescence experiment, whereas red arrows indicate non-transfected control cells. (i) Quantification of filopodia number per cell and filopodia length in control cells, FLOT2-expressing and IRSp53(4K)-expressing cells. Scale bars are indicated in each panel.

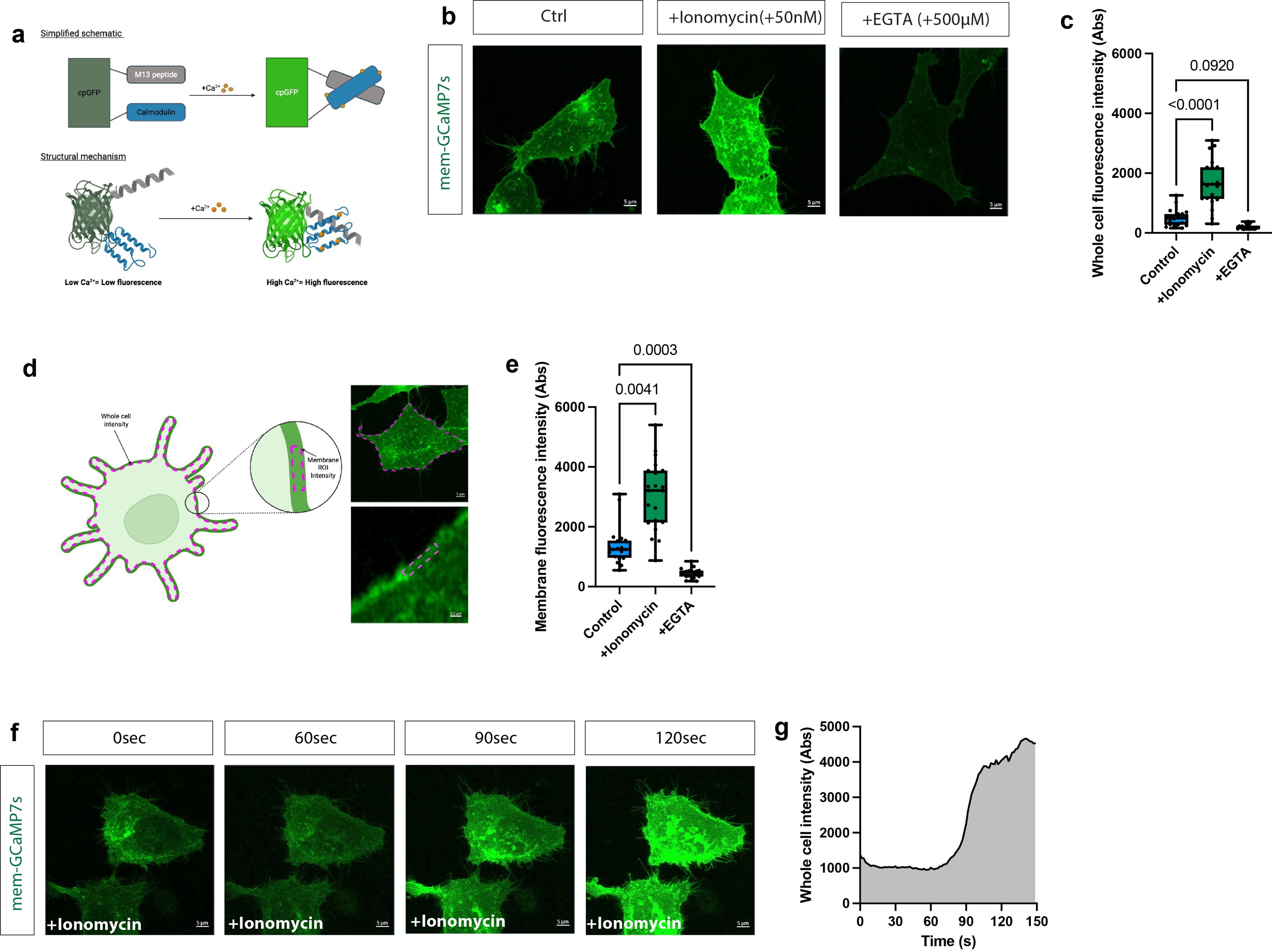

**Supplementary Figure 2. Functional validation of the membrane-targeted GCaMP7s calcium reporter in HeLa cells.** (a) Schematic showing the principle of GCaMP7s fluorescence activation by calcium binding. Calcium binding induces a conformational change in the cpGFP–calmodulin sensor, increasing fluorescence intensity. (b,c) HeLa cells expressing membrane-targeted GCaMP7s were treated with ionomycin or EGTA to validate reporter responsiveness. Representative images and whole-cell fluorescence quantification show increased GCaMP7s fluorescence after ionomycin treatment and reduced signal after EGTA treatment. (d,e) Strategy for membrane-associated GCaMP7s quantification. Regions of interest were drawn along the cell membrane and filopodia to quantify membrane-localised calcium reporter fluorescence. Membrane fluorescence intensity increases after ionomycin treatment and decreases after EGTA treatment. (f,g) Time-lapse imaging of membrane-GCaMP7s after ionomycin addition confirms rapid reporter activation. Representative images show increasing fluorescence over time, and quantification shows whole-cell fluorescence intensity during the imaging period. GCaMP7s signal is shown as raw fluorescence intensity. Data are presented from multiple cells per condition. Statistical significance was determined using the indicated multiple-comparison test; *p*-values are shown in the graphs. Scale bars are indicated in each panel.

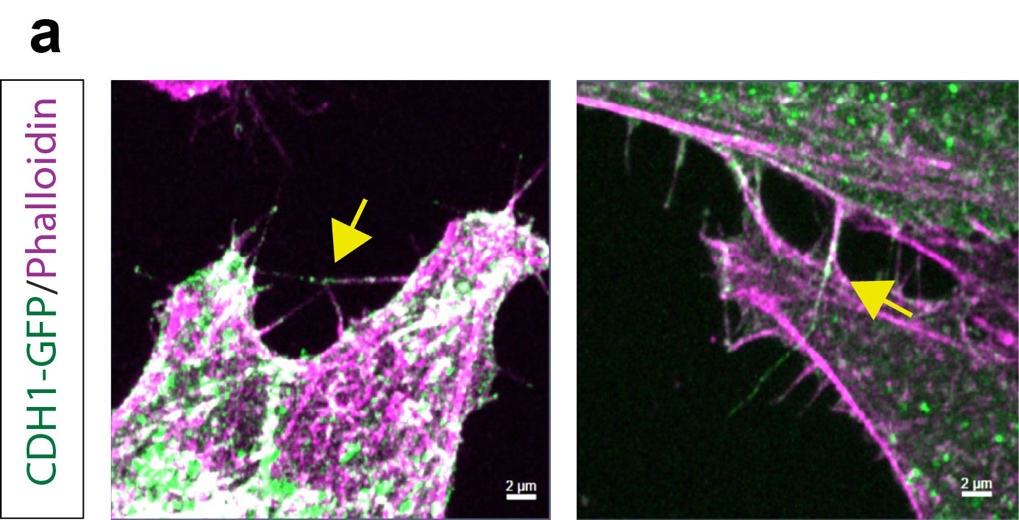

**Supplementary Figure 3. Localisation of CDH1-GFP after overexpression in HeLa cells.** (a) yellow arrows indicate accumulation of CDH1 at cytoneme contact sites.

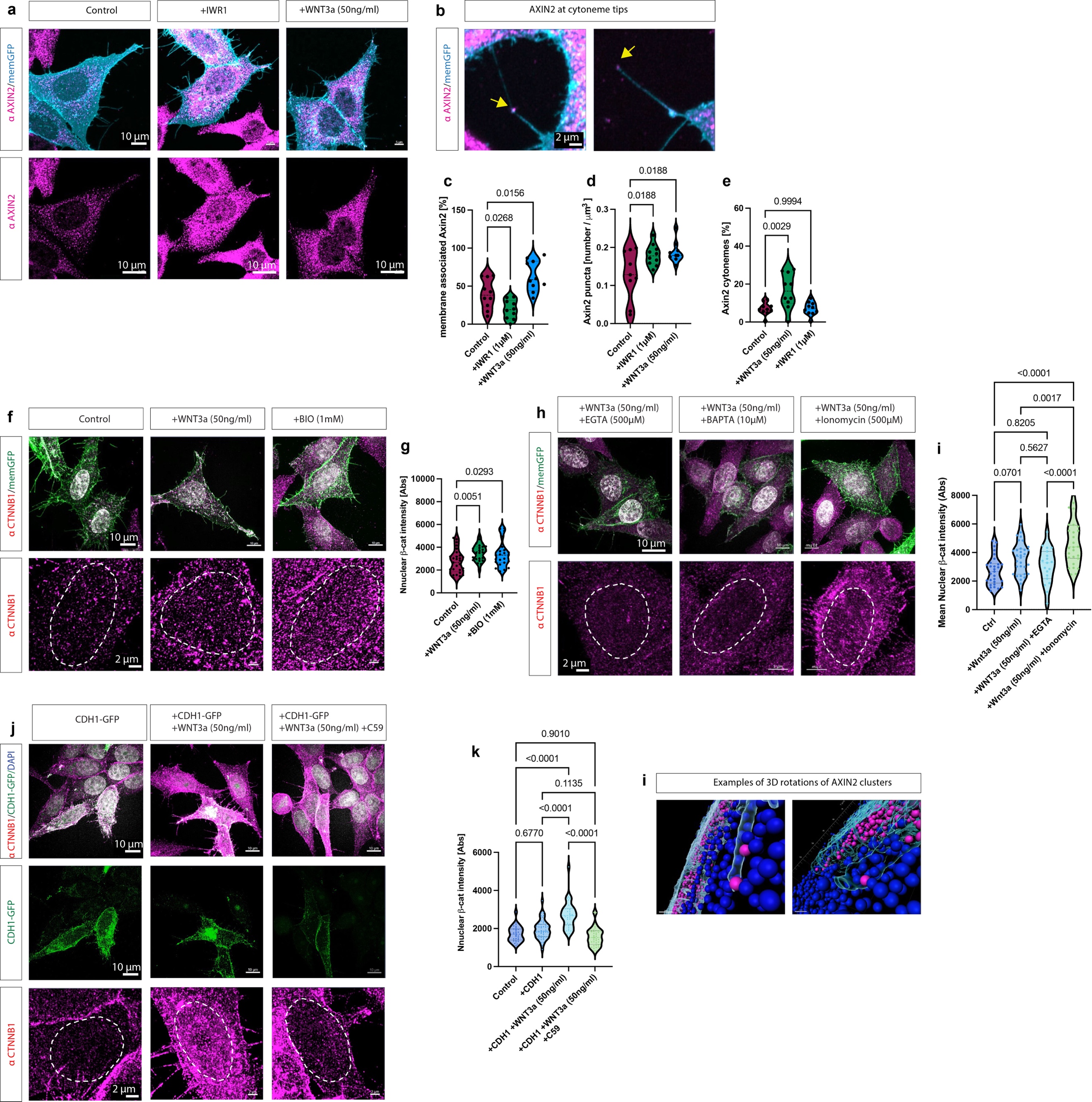

**Supplementary Figure 4. Pharmacological validation of AXIN2 recruitment at cytonemes and β-catenin accumulation in the nucleus .** (a–e) AXIN2 localisation was analysed in membrane-GFP-expressing HeLa cells under control conditions, after IWR1 treatment, or after recombinant WNT3a stimulation. Representative images show total and membrane-associated AXIN2 staining. Higher-magnification images highlight AXIN2-positive puncta at cytoneme tips. Quantifications show membrane-associated AXIN2, AXIN2 puncta density and the percentage of AXIN2-positive cytonemes. Yellow arrows indicate AXIN2-positive cytoneme tips. Nuclear β-catenin/CTNNB1 localisation was analysed as a downstream readout of canonical Wnt pathway activation after treatment with recombinant WNT3a protein and the GSK3 inhibitor BIO1 (f,g) and Calcium modulators EGTA, BAPTA, and ionomycin (h, i). CDH1-GFP expression in combination with WNT3a treatment alters CTNNB1 accumulation (j, k). Nuclear β-catenin was quantified as mean nuclear CTNNB1 fluorescence intensity; dashed outlines indicate nuclei. Data are presented from multiple cells per condition. Statistical significance was determined using the indicated multiple-comparison test; *p*-values are shown in the graphs. Scale bars are indicated in each panel. (i) Display of 3D rotations after surface rendering to determine AXIN2 clusters in close proximity to cytoneme contacts.
